# Advancing Liver Cancer Precision Medicine with TARGET-SL

**DOI:** 10.64898/2026.05.19.725819

**Authors:** Rhys Gillman, Benjamin J. Dwyer, Sara Pasic, Gayatri D Shirolkar, Nathan Main, The Liver Cancer Collaborative, Matt A. Field, Ulf Schmitz, Lionel Hebbard

**Affiliations:** Department of Physical Sciences, College of Science and Engineering, James Cook University, Townsville, Queensland, Australia; Centre for Tropical Bioinformatics and Molecular Biology, James Cook University, Cairns, Queensland, Australia; Curtin Medical School and Curtin Medical Research Institute, Curtin University, Bentley, WA 6102, Australia; Harry Perkins Institute of Medical Research and Centre for Medical Research, University of Western Australia, 35 Stirling Highway, Perth WA 6009, Australia; Immunogenomics Lab, Garvan Institute of Medical Research, Darlinghurst, New South Wales, Australia; Menzies School of Health Research, Charles Darwin University, Darwin, Northern Territory, Australia; Faculty of Medicine & Health, The University of Sydney, Camperdown, Australia; Department of Biomedical Sciences and Molecular and Cell Biology, College of Medicine and Dentistry, College of Science and Engineering, James Cook University, Townsville, Queensland, Australia; Storr Liver Centre, Westmead Institute for Medical Research, Westmead Hospital and University of Sydney, Sydney, New South Wales, Australia; Australian Institute for Tropical Health and Medicine, Townsville, Queensland, Australia

**Keywords:** Driver gene prediction, drug prediction, bioinformatics, precision medicine, primary liver cancer

## Abstract

**Background and Aims:** A major goal of personalised liver oncology is the ability to make targeted predictions about cancer-specific toxicity, however there are limited methods available. To address this, we validated the performance of our bioinformatics framework, TARGET-SL, through *ex vivo* drug screening.

**Methods:** Using TARGET-SL we predicted gain of function (GOF), loss of function (LOF) and synthetic lethal (SL) genetic events, and corresponding drug candidates. We validated drug predictions across hepatocellular carcinoma (HCC) cell lines, and a cohort of HCC and cholangiocarcinoma (CCA) patient-derived organoids (PDOs).

**Results:** For HCC cells and PDOs we found 37.5% and 25% of the respective selected compounds induced unique target-specific growth inhibition based on genetic biomarkers, suggesting novel biomarker-driven drug sensitivities.

**Conclusions:** Our analyses demonstrate TARGET-SL’s potential to enhance personalized drug screening for liver cancer, by focusing on genetically informed targets. This will reduce experimental costs and accelerate the pace of therapeutic discovery.

**Impact and Implications:** Primary liver cancer (PLC) is a cancer with poor prognosis, and current therapies increase survival only for a minority of patients. Through the application of TARGET-SL we can predict, for each patient, the essential genes and corresponding small molecule inhibitors. These data support further investigation in larger patient cohorts and offer the possibility to specify new small molecule inhibitors and to repurpose current drugs for PLC treatment.

**Graphical Abstract:** 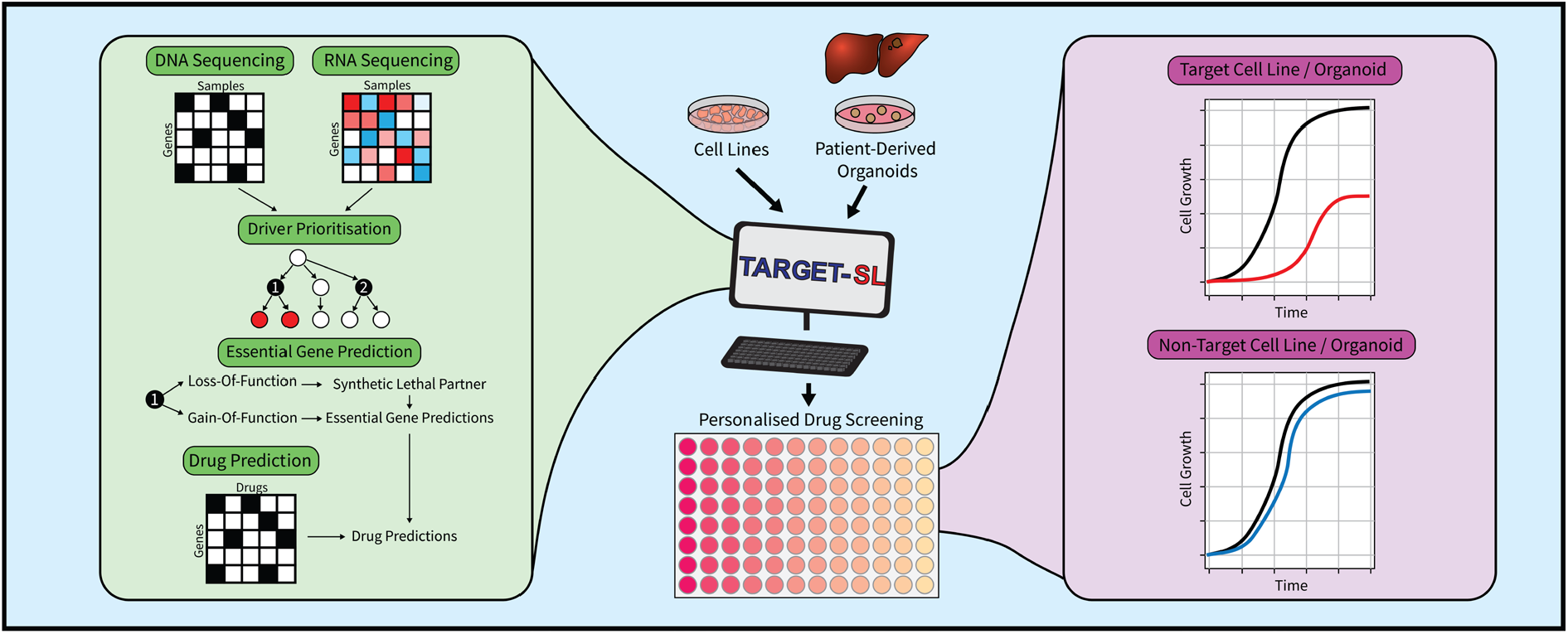

**Highlights:**

- TARGET-SL can predict gene and drug sensitivities for cell lines and patient-derived organoids
- This may reduce drug screening costs and accelerate the pace of therapeutic discovery.
- TARGET-SL may assist in the repurposing of current drugs and their rapid translation for primary liver cancer
- TARGET-SL is tumour-type agnostic, and therefore may have application in other cancers with poor prognosis

## Introduction

Primary liver cancer (PLC), which includes hepatocellular carcinoma (HCC) and cholangiocarcinoma (CCA), is the sixth most common cancer worldwide and the third leading cause of cancer-related deaths [1]. PLC associates with poor outcomes, and five-year survival rates near 20% [2]. Moreover, the limited therapeutic options available offer only marginal improvements to patient survival, are effective in just 30% of patients, and can, in some instances, promote tumour progression and adverse effects [3, 4].

The limited efficacy of these therapeutics may be attributable to PLC having diverse genetics, with a long tail of infrequent mutations across patients [5]. Notwithstanding, patients are commonly treated without genetic information. Thus, there is an urgent need to personalise liver cancer oncology, by identifying drugs with tumour-specific toxicity.

In other cancers, *ex vivo* drug screening platforms are emerging as a promising approach to identify such drugs [6]. These are often performed in high-throughput assays, involving hundreds to thousands of drugs per patient, yet they achieve hit rates of just 0.5-5% [7-9]. Additionally, such screens are costly, making them less clinically viable.

To address these issues, algorithmic approaches predicting patient-specific drug sensitivities can help narrow the search space for personalised screenings of targeted therapies. Hence, we developed <u>T</u>umour-specific <u>A</u>lgorithm for <u>R</u>anking <u>GE</u>netic <u>T</u>argets via <u>S</u>ynthetic <u>L</u>ethality (TARGET-SL) [10], that predicts essential genes, synthetic-lethal (SL) interactions and corresponding small molecule inhibitors for individual patients. Here, we demonstrate TARGET-SL’s utility by validating predictions in HCC cell lines and a cohort of HCC and CCA patient-derived organoids (PDOs).

## Methods

The materials and methods used are described in the supplementary information.

## Results

### Validation of TARGET-SL Predictions in Hepatocellular Carcinoma Cells

We used TARGET-SL to predict essential genes and drug sensitivities in 1290 cell lines using publicly available Cancer Cell Line Encyclopaedia (CCLE) data [10, 11]. We selected 8 drug predictions for four HCC cell lines, Huh7, Hep3B, SNU-423 and PLC/PRF/5 for *in vitro* validation (**Supplementary Table S1**). The anticipated effects of these predictions are based on high-throughput CRISPR-KO screening and/or small-molecule drug screening from the CCLE, and are shown in (**Supplementary Figure 1**).

We also screened with Sorafenib, a common HCC drug. At the lowest concentration, 5uM Sorafenib slowed the growth of immortalised healthy hepatocyte (IHH) cells after 72hrs, while the effects on cancer cell lines were limited. At higher concentrations, Sorafenib significantly impeded growth in both cancer and IHH cells, reflecting non-cancer-specific toxicity (**Figure 1A and B**).

**Figure 1:**
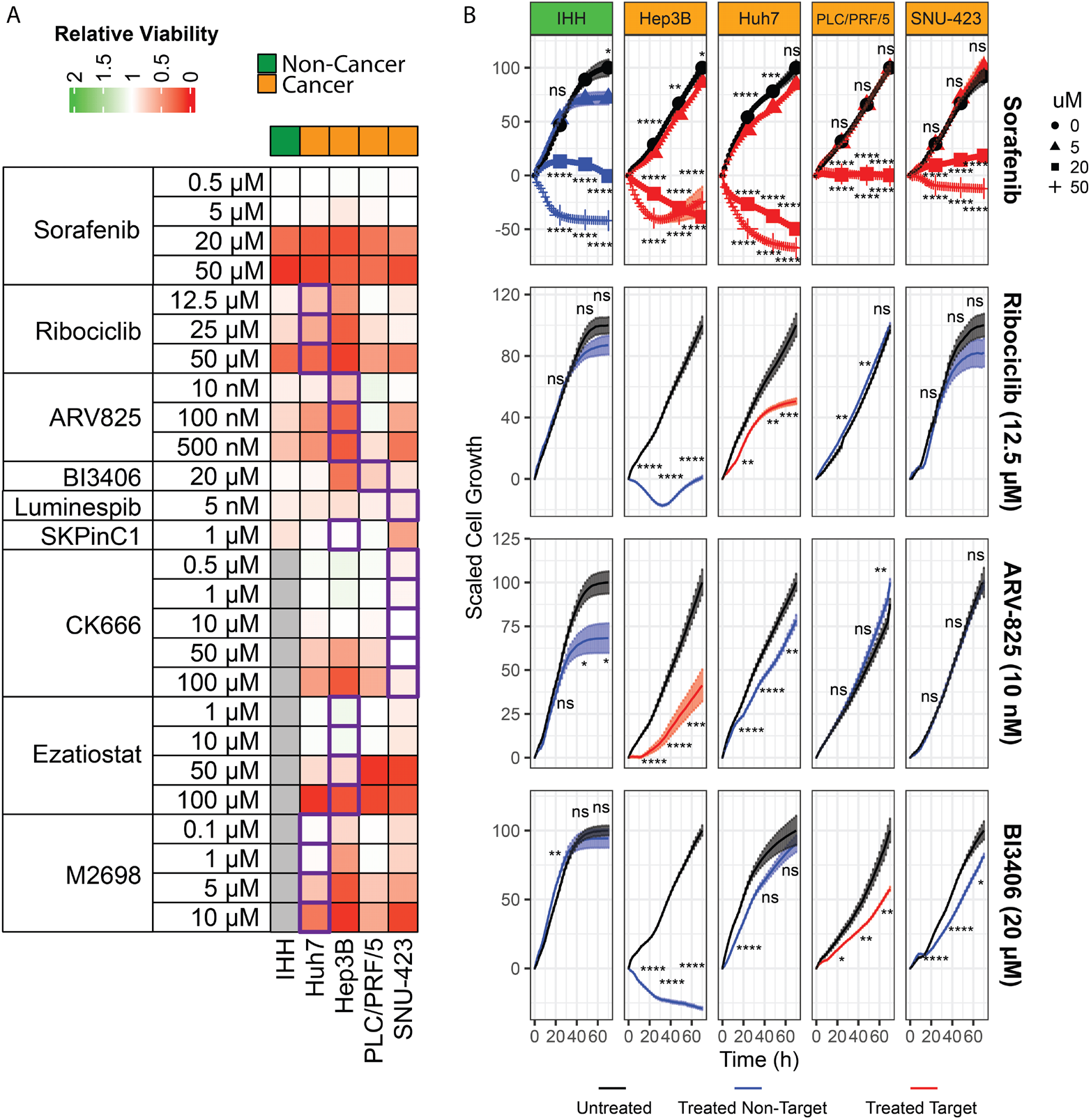
Validation of TARGET-SL drug predictions *in vitro*. The HCC cell lines Hep3B, Huh7, PLC/PRF/5, and SNU-423, and immortalised healthy hepatocytes (IHH) were exposed to Sorafenib and eight TARGET-SL-predicted drugs at various concentrations. (A) Heatmap of the average Relative Viability of each cell line treated after 72 hours. Purple boxes indicate the target cell line of each TARGET-SL-predicted drug. Grey boxes indicate missing data. (B) Selected growth curves of TARGET-SL-predicted drugs showing target-specific toxicity. Cell confluence was calculated and scaled to starting and maximal confluence per cell line (n=8-12). Significance at 24, 48, and 72 hours was determined using T-tests with Benjamini-Hochberg correction.

We observed direct cancer- and target-specific toxicity in Huh7, Hep3B, and PLC/PRF/5 cells following treatment with their respective TARGET-SL-predicted drugs, Ribociclib, ARV-825 and BI3406 (**Figure 1A and B**), representing 3 out of 8 (37.5%) drugs screened. Critically, these three drugs showed limited toxicity to the control IHH cell line. In some cases, the non-target cancer cells were affected by the screened drugs, including Hep3B sensitivity to Ribociclib and BI3406, and Huh7 sensitivity to ARV-825. Luminespib and SKPinC1 showed universal toxicity across all cell lines and/or higher sensitivity in IHH cells than tumour cells and were not investigated further.

Unexpectedly, 3/8 (37.5%) of the predicted drug sensitivities showed cell-specific drug resistance, including CK666, Ezatiostat, and M2698 (**Figure 1A and Supplementary Figure 2A**). To test whether this was due to incorrect loss or gain-of-function annotation, we treated the SNU-423 *ACTR3* mutation as a loss-of function (LOF) and targeted its putative SL-partner, KRAS, with Salirasib. Indeed, even though Salirasib showed some toxicity to IHH cells, SNU-423 cells had no resistance to Salirasib (**Supplementary Figure 2B**).

### Validation of TARGET-SL Predictions in Patient-Derived Organoids

To demonstrate the *ex vivo* applicability of TARGET-SL for patient samples, we made drug predictions using tumour sequencing data from three HCC and three CCA patients with matched PDOs for drug sensitivity validation, combined with TCGA data (**Supplementary Figure 3**) to allow calculation of differential gene expression. We generated and manually curated a list of eight drug predictions for the patients based on high-ranking TARGET-SL predictions and availability of targeted small-molecule inhibitors (**Supplementary Table S2**).

Drugs were screened against the patient-matched cancer PDO lines, and human hepatocytes derived from humanised liver FRG mice (hFRG-Heps) [12] as a non-tumour control, over 4 days. Two drugs, AZD5582 and Mithramycin A showed significant target-specific toxicity (**Figure 2**). AZD5582 also showed off-target toxicity to PCB000314, but minimal toxicity to hFRG-Heps, while Mithramycin A reduced viability in all PDOs but to a lesser extent in hFRG-Heps. C25-140 showed the greatest toxicity in the target PDO PCB000087, though the effect was non-significant, suggesting that higher concentrations are required to elicit an effect. SCH-900776 impaired the growth of its target PDO, PCB000087, more than other cancer PDOs, but was toxic to hFRG-Heps, and consequently not cancer-specific. Finally, in agreement with our cell line results, we observed that some TARGET-SL drug sensitivity predictions in fact translated to drug resistant profiles, most notably GSK-J4 in both PCB000166 and PCB000061 (**Figure 2A**).

**Figure 2:**
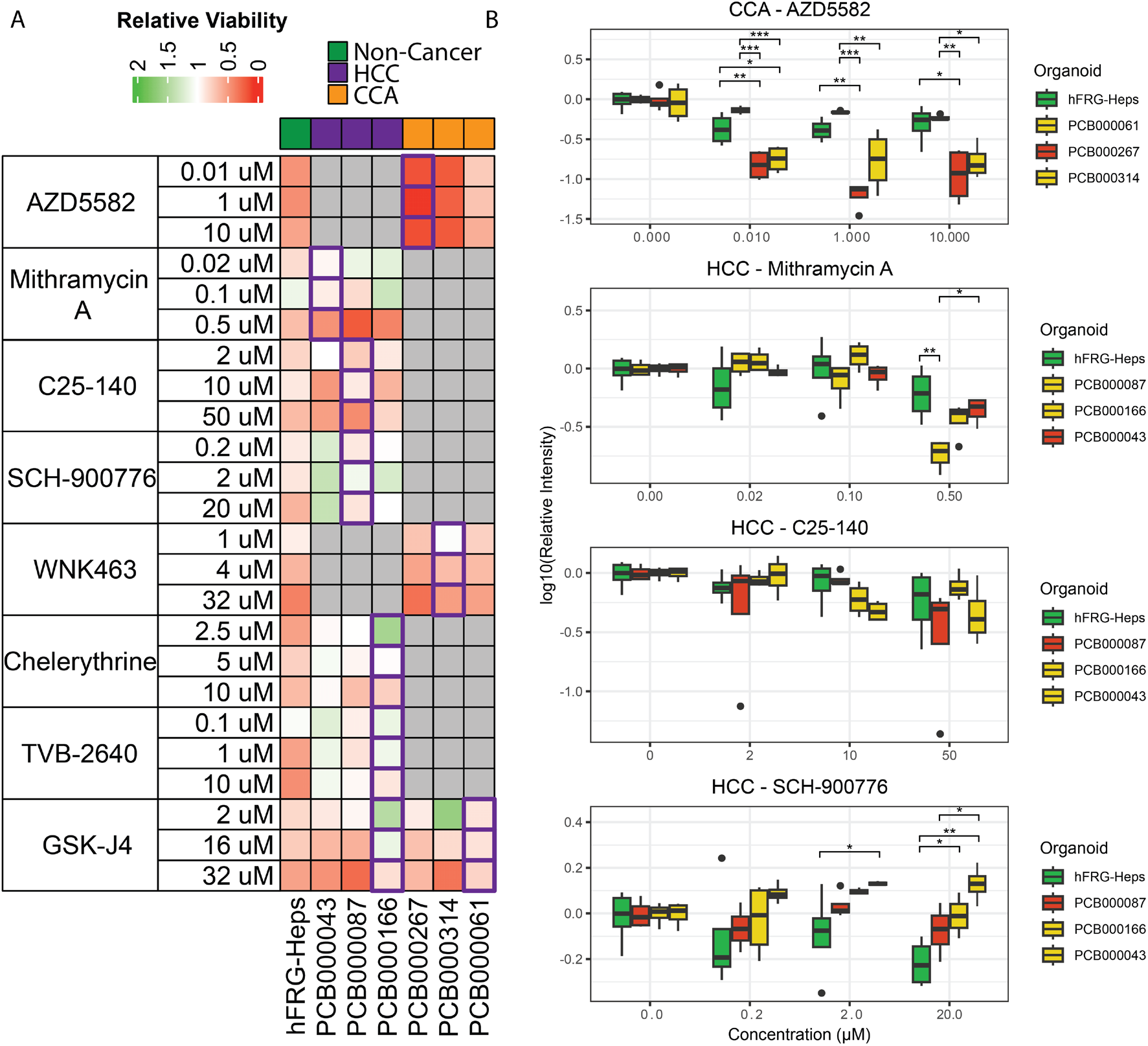
Validation of PDO TARGET-SL drug predictions *ex vivo*. TARGET-SL drug predictions were made using sequencing data from patient cancer tissue with matched PDOs, including three HCC patients (PCB000043, PCB000087, PCB000166), three CCA patients (PCB000267, PCB000314, PCB000061), and hFRG-Heps a non-cancer control. Relative viability (relative to DMSO control) was assessed through a Cell-Titre Glo 3D Assay. (A) Heatmap of average Relative Viability of each PDO treated. Purple boxes indicate the target PDO of each TARGET-SL-predicted drug. (B) Selected relative viabilities of drugs showing target-specific PDO toxicity. Red boxes indicate target cancer organoids, yellow boxes indicate non-target cancer organoids, green boxes indicate control organoids. Significant differences were determined using ANOVA with Tukey Honestly Significant Differences.

## Discussion

It is expected that *ex vivo* drug screening will become an increasingly important clinical component for future oncology. However, the complexity of targeted therapies in a landscape of extensive tumour genetic heterogeneity means that approaches that reduce the search space of drug candidates are urgently required. Here we applied TARGET-SL to predict targeted drug candidates and tested these *ex vivo* using HCC cell lines and PLC patient-derived organoids.

We showed that three drugs, Ribociclib, ARV-825, and BI-3406 demonstrated *in vitro* target-specific cell toxicity, which was consistent with our TARGET-SL *in silico* drug screening results [10]. Notably, each of these drugs or their gene targets have shown therapeutic potential in previous studies. However, none of these previous studies considered the sensitisation of target cells due to the presence of specific mutations. TARGET-SL predicted Ribociclib, a CDK4 inhibitor [13], to be effective in Huh7 cells, where CKD4 was identified as an essential gene due to its putative SL-interaction with HDAC1, a gene mutated in these cells. Interestingly, CDK4 inhibitors have been studied in HCC, though not in the context of HDAC1 SL [14], and the combined inhibition of CDK4 and HDAC1 has shown efficacy *in vitro* for melanoma treatment [15].

ARV-825 inhibits BRD4 and was predicted to be essential in Hep3B cells based on a SL-relationship with *RB1*. Hep3B is one of the few HCC cell lines that has an *RB1* mutation [16]. Our results show Hep3B-sensitivity to ARV-825, which is supported by another study, illustrating that Hep3B cells are more sensitive to BRD4 inhibition by the inhibitor JQ1, than other HCC cell lines [17]. Hence, TARGET-SL explains that this sensitisation in Hep3B cells is due to a mutation in the BRD4 SL-partner, *RB1*.

We found that a GOF variant in PLC/PRF/5 cells sensitises these cells to the SOS1 inhibitor BI-3406. SOS1 is not well studied in HCC, however it is frequently over-expressed and regulates epithelial-to-mesenchymal transition and invasion in cell line models [18]. Together, these results suggest that TARGET-SL can improve the application of previously studied drugs through identifying personalised genetic variants that sensitise a tumour to a drug.

In some cases, drug predictions unexpectedly showed resistance in their target cell line rather than sensitivity, namely CK666, Ezatiostat, and M2698 (**Supplementary Figure 2A**). This could be explained as mutations being incorrectly assigned as LOF or GOF mutations, which we have discussed [10]. We investigated this by targeting KRAS, a SL-partner of the target gene, *ACTR3* in SNU-423 cells, and indeed found these cells were sensitive to the drug (**Supplementary Figure 2B**), suggesting that *ACTR3* mutations in SNU-423 cause LOF. This suggests a limitation in LOF and GOF assignment by TARGET-SL, a component which we anticipate will improve over time, with improvements in experimental and prediction databases that TARGET-SL utilises.

To demonstrate a proof of principle for targeted *ex vivo* screening we applied TARGET-SL to HCC and CCA patient PDOs. The best performing drug in this cohort was AZD5582, which targets a GOF mutation in *BIRC3* in a CCA organoid. BIRC3 is a potential prognostic biomarker in HCC and associates with cancer growth in cell lines and mouse models through siRNA [19] and drug inhibition [20]. Our results also suggest potential efficacy of targeting SP3 with mithramycin A in our *ex vivo* PDO model. Mithramycin A has been considered for cancer treatments, but its therapeutic dose is limited by the development of hepatotoxicity [21]. Our results suggest that SP3 GOF mutations could sensitise cancer cells to mithramycin A, which could lead to lowering the required therapeutic dose to reduce hepatoxicity.

This validation study identified only a small number of cancer-specific agents, we nonetheless demonstrate a significantly higher success rate for HCC cells (37.5 %) and PDOs (25 %), when compared with expensive high-throughput drug screening studies, which usually lie at < 5% [7-9]. This, therefore, demonstrates the power of TARGET-SL for performing targeted screens rather than high-throughput screening, and offers a small step in the direction of developing precision medicine for PLC patients.

In conclusion, we demonstrated that TARGET-SL can predict target-specific PLC cancer sensitivity to small molecule inhibitors based on the prediction of cancer essential genes, having the potential to improve the hit rate of *ex vivo* drug screening.

## The Liver Cancer Collaborative Contributors

Individual contributors to the The Liver Cancer Collaborative, with specific affiliations in addition to those listed for the main authors are:

Peter Leedman^4^

Nina Tirnitz-Parker^3^

Louise Winteringham^4^

Michael C. Wallace^11,12^

Jonathan Tibballs^13^

Benjamin Dwyer^3^

Gayatri Shirolkar^3^

Ashlin Donnellan^4^

^11^Department of Hepatology, Sir Charles Gairdner Hospital, Perth, WA, Australia.

^12^Medical School, University of Western Australia, 35 Stirling Highway, Perth WA 6009, Australia

^13^Medical Imaging Department, Sir Charles Gairdner Hospital, Perth, WA, Australia.

## Ethics Statement

This research has been conducted using samples and data from The Perkins Cancer Biobank. The Biobank has been approved by WA Health Central Human Research Ethics Committee approval number RGS0000000919. Patient-derived organoids are used under Curtin ethics approval HRE2021-0203.

