## Supplementary Methods for "Advancing Liver Cancer Precision Medicine with TARGET-SL"

*TARGET-SL Cell Line Drug Predictions*

TARGET-SL [1] was previously used to generate drug predictions for 1290 cell lines in the Cancer Cell Line Encyclopaedia (CCLE) [2]. The top drug predictions for cell lines Huh7, Hep3B, PLC/PRF/5, and SNU-423 were manually curated, and drugs were selected based on literature support and purchasing availability. These included M2698 (Cayman Chemical, #36816), Salirasib (Cayman Chemical, #10010501), Ezatiostat (Cayman Chemical, #16248), CK-666 (Cayman Chemical, #29038), LEE011 (Cayman Chemical, #17666), BI-3406 (Cayman Chemical, #35261), SKPin C1 (Selleck Chemicals, #S8652), and ARV-825 (Selleck Chemicals, #S8297).

*Cell Lines*

Huh7 and Hep3B cells were cultured in Dulbecco's Modified Eagle's Medium (DMEM) high glucose supplemented with 10% foetal bovine serum (FBS) and 1% penicillin/streptomycin (P/S). SNU-423 and PLC/PRF/5 cells were cultured in RPMI-1640 medium supplemented with 10% FBS, 1% P/S and 2mM L-glutamine. IHH cells were cultured in DMEM with 10% FBS, 1% P/S, 50 nM dexamethasone, 20 mU/mL insulin (all cell culture reagents sourced from Merck). All cell lines were cultured at 37°C with 5% $CO_2$, and media was refreshed every 3 days.

*Cell Line Drug Screening*

Drugs were initially screened across the four cancer cell lines to determine the best screening concentration and drugs with the most variable sensitivity. Cells were seeded overnight into 96-well plates at 5000 cells per well in replicates of 3 per concentration, and their medium replaced with fresh drug-containing medium in <0.5% DMSO. Cell confluence was monitored for 3-4 days using an Incucyte SX1 live-cell imaging system (Sartorius). Drugs which demonstrated target-specific toxicity were subjected to an additional screen, which included IHH cells and performed using 8 replicates at each drug concentration.

*Cell Line Screening Analysis*

Confluence values were scaled between each cell line's starting confluence and maximum confluence to normalise across cells with differing growth rates and morphology. Growth of untreated and treated cells were compared at 24, 48 and 72 hours using a Student's T Test with Benjamini-Hochberg Correction.

*Patients*

We included three HCC patients and three CCA patients in our study with patient tissue, sequencing data and PDOs provided by The Liver Cancer Collaborative and the Perkins Cancer Biobank. The Biobank has been approved by WA Health Central Human Research Ethics Committee approval number **RGS0000000919.** The PDOs were generated by Dr Ben Dwyer.

*Patient Sequencing*

RNA sequencing (RNA-seq) (Illumina NovaSeq) and whole-exome sequencing (WES) (Illumina Novaseq) was carried out by GenomicsWA. RNA sequencing reads were aligned to the human reference genome (hg38, Gencode v32) using STAR (v.2.7.5a) [3], and gene-level read counts were quantified using featureCounts (v2.0.1) [4]. WES sequencing reads were aligned to the human reference genome using Burrows-Wheeler Aligner (BWA, v0.7.17) [5] and processed following Genome Analysis Toolkit (GATK) best-practices [6]. Variants were called using Mutect2, Varscan

2 [7], MuSE [8], and Strelka [9]. Variant annotation was performed using Funcotator (v1.7). Copy number variants were called using CNVkit [10].

*Sequencing Analysis*

For RNA-seq analysis, gene-wise count matrices were merged with gene counts from The Cancer Genome Atlas (TCGA) [11], including 50 normal and 373 HCC cancer samples, and 9 normal and 36 CCA cancer samples. Count data were variance-stabilised using DESeq2 [12] and batch-corrected using Limma's removeBatchEffect function in R [13]. We assessed the quality of the batch correction using principal component analyses (PCA) based on the top 3000 most variable genes (**Supplementary Figure 3**). Similarly, we retrieved somatic non-synonymous coding mutations from TCGA samples, lifting them over to hg38 using Liftover [14], and gene-wise log2 copy number (Log2CN) values.

TARGET-SL was used to predict essential genes and drug sensitivity for each LCC patient, using the TCGA as additional samples for calculating differentially-expressed genes. For personalised driver prioritisation, four algorithms were applied: DawnRank [15], OncoImpact [16], PersonaDrive [17], and sysSVM2 [18].

*Organoid Drug Screening*

Organoids were generated from fresh biopsy or resection tissue from each patient. 384-well plates were prepared with 10 µL of Cultrex Reduced Growth Factor BME-2 (R&D Systems) and incubated overnight at 4°C. Prior to solidification, plates were centrifuged at 1000 x g to flatten BME-2, then incubated at 37°C for 30 min. Organoids were dissociated to single cells by incubation with trypsin-EDTA for 5 min. Cells were counted using a haemocytometer, centrifuged at 300 x g for 5 min and resuspended at 10,000 cells/ml in 40 ml organoid medium (Advanced DMEM F12 (Thermofisher Scientific), 1% P/S (Thermofisher Scientific), Glutamax (Thermofisher Scientific), 10 mM HEPES pH 7 (Sigma-Aldrich), 1 x N2 supplement (Thermofisher Scientific), 1 x B27 supplement (Thermofisher Scientific), 10 mM Nicotinamide (Sigma-Aldrich), 50 ng/mL EGF (Stem Cell Technologies), 10 nM Gastrin (Sigma-Aldrich), 100 ng/mL FGF10 (Thermofisher Scientific), 25 ng/mL HGF (Stem Cell Technologies), 500 ng/mL R-Spondin 1 (Stem Cell Technologies), 1.25 mM N-Acetylcysteine (Sigma-Aldrich), 5 µM A-83-01 (Sigma-Aldrich), 10µM Forskolin (Stem Cell Technologies) [19]. For drug treatment, cells were seeded into 384 well plates containing warmed BME-2 at 1000 cells/well by dispensing 100µl/well of cell suspension using a ViaFlo384 automated liquid handling system. Cells were cultured at 37°C with 5% $CO_2$ for 7 days to allow organoids to develop, after which the medium was replaced with 50µL of fresh organoid medium and supplemented with 50µL of organoid medium containing 2 times the final concentration of each drug treatment in quadruplicate. DMSO concentration was maintained at 0.2% for all drug treatments. Organoids were cultured for an additional 4 days after exposure to drug treatments.

*Organoid Viability*

Total organoid viability was assessed using a Cell Titre Glo® 3D assay (Promega), according to manufacturer instructions. The relative intensity was calculated by dividing the intensity of each treatment well by the mean of DMSO-control wells for each organoid line. Comparisons were made between organoids at each drug concentration using ANOVA with Tukey Honestly Significant Differences test.
