## Supplementary Figures for "Advancing Liver Cancer Precision Medicine with TARGET-SL"

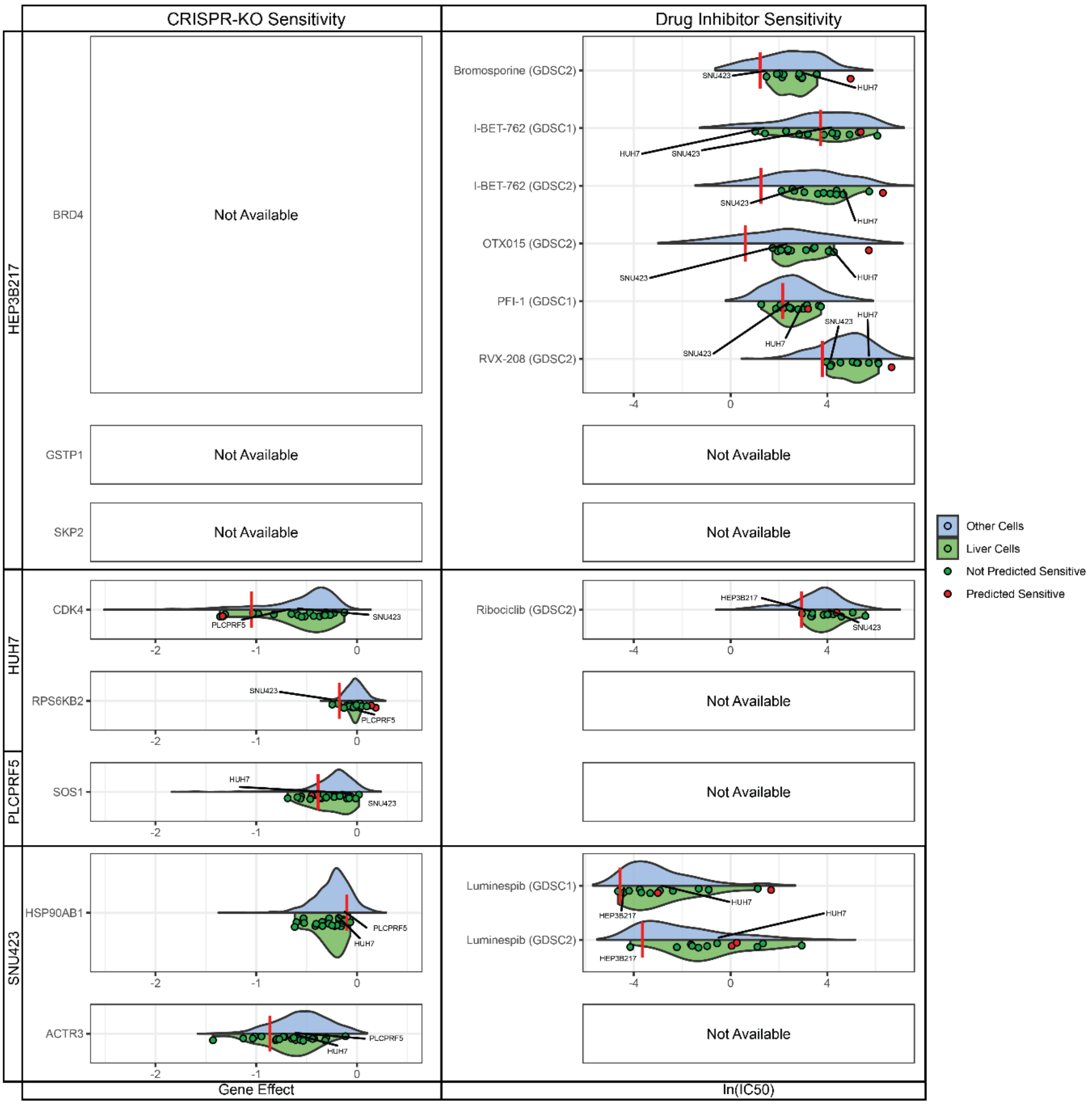


**Supplementary Figure 1.** Cancer Cell Line Encyclopedia (CCLE)-based validation of TARGET-SL predictions. CRISPR-KO gene effect (Left Panel) from DepMap shows the effect of specific gene knockouts on each cell line, where more negative values indicate less cell growth. The ln($IC_{50}$) (right panel) shows the effect of small-molecule inhibitors of each gene from the left panel on the growth of each cell line. All cells in the CCLE were examined (n=1290), and divided into the target cell line (red line), cell lines of the same cell type predicted to be sensitive to gene-KO or drug (red points), cell lines of the same cell type not predicted to be sensitive (green points), the distribution of values from cells of the same cell type (green violin), and all other cells (blue violin).

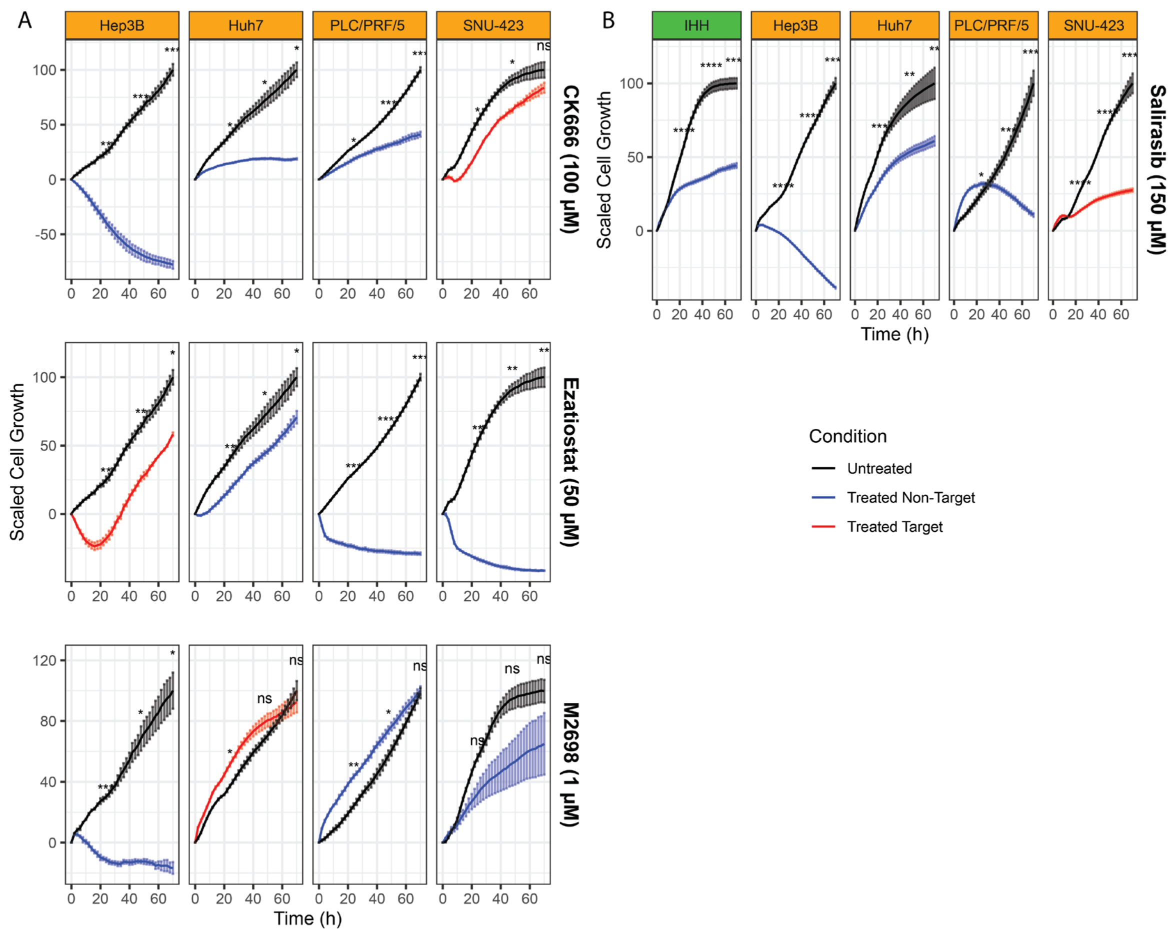

**Supplementary Figure 2**: TARGET-SL-predicted drugs showing target-specific resistance. (A) Growth curves showing three drugs with marked resistance in their target cell lines, CK666, Ezatiostat, and M2698 (n=3). (B) SNU-423 cells were screened using Salirasib as an alternative drug (n=8). Cell confluence was calculated and scaled to starting and maximal confluence per cell line. Significance at 24, 48, and 72 hours was determined using T-Tests with Benjaminni-Hochberg Correction.

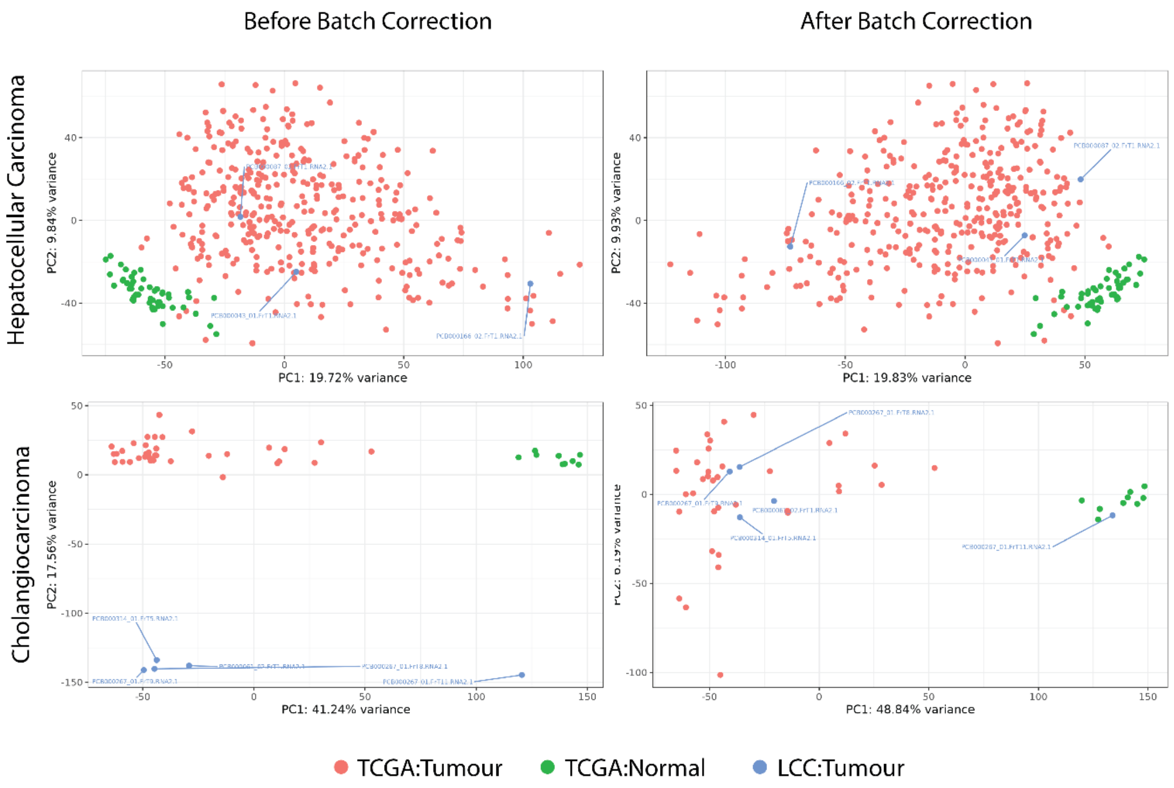


**Supplementary Figure 3**: PCA plots of the top 3000 most-variable genes across all samples before and after batch correction. Plots are divided into hepatocellular carcinoma and cholangiocarcinoma cohorts, including tumour and normal samples from The Cancer Genome Atlas (TCGA) and patient tissue samples from the Liver Cancer Collaborative (LCC), from which the PDOs were used for *ex vivo* studies. Note that three replicate samples were available for a single cholangiocarcinoma patient, PCB000267. The sample PCB000267_01.FrT11.RNA2.1 clustered with normal tissue samples, suggesting non-cancer contamination, and was removed, while the other two samples were closely clustered and were therefore combined by taking their average expression for all genes.
